# Simultaneous profiling of chromatin accessibility and methylation on human cell lines with nanopore sequencing

**DOI:** 10.1101/504993

**Authors:** Isac Lee, Roham Razaghi, Timothy Gilpatrick, Michael Molnar, Norah Sadowski, Jared T. Simpson, Fritz J. Sedlazeck, Winston Timp

## Abstract

Understanding how the genome and the epigenome work together to control gene transcription has applications in our understanding of diseases such as human cancer. In this study, we combine the ability of NOMe-seq to simultaneously evaluate CpG methylation and chromatin accessibility, with long-read nanopore sequencing technology, a method we call nanoNOMe. We generated >60Gb whole-genome nanopore sequencing data for each of four human cell lines (GM12878, MCF-10A, MCF-7, MDA-MB-231) including repetitive regions inaccessible by short read sequencing. Using the long reads, we find that we can observe phased methylation and chromatin accessibility, large scale pattern changes, and genetic changes such as structural variations from a single assay.

## INTRODUCTION

Many human diseases, including all forms of neoplasia, result from aberrant gene regulation through mechanisms including genetic mutation, altered signal transduction, and epigenetic alteration. Healthy cells use tight control of the epigenome to modulate active transcription of genes through the coordination of numerous signals including methylation of CpG dinucleotides, chromatin accessibility, and nuclear organization. The epigenome is highly mutable, changing dynamically in response to external stimuli, which can result in epigenetic variation among phenotypically and genetically homogeneous populations. This feature is especially evident when comparing tissue-paired normal and cancer samples^1^. Normal tissue commonly has well-defined epigenetic signatures, in contrast to transformed cells wherein cancer epigenetics is much more varied from sample to sample^1^, and even from cell to cell^2^.

With the proliferation of DNA sequencing technologies, methods have been developed for examining nuclear organization, protein binding site occupancy, chromatin accessibility, and methylation state. Many of these methods rely on the vulnerability of accessible chromatin to enzymatic treatment, e.g. DNAse-seq^3^, ATAC-seq^4^. One of these methods, NOMe-seq^5^, labels genomic regions in a nucleosome-depleted, accessible state using an exogenous GpC methyltransferase. Combined with bisulfite conversion, this method permits simultaneous evaluation of the endogenous cytosine methylation as well as nucleosome occupancy. *However, these methods do not directly interrogate the DNA strand, and the reads are typically too short to provide information about the regional context of the DNA.*

The long reads possible with nanopore sequencing provide a deeper level of insight, allowing investigation of long-range patterns on individual DNA molecules. We and others have previously shown that endogenous CpG methylation can be accurately called with nanopore data^6,7^. By extending this model to include the non-native GpC modifications, we are able to adapt the NOMe-seq workflow to the nanopore platform. We then take advantage of the long read lengths (>10kb) generated by nanopore sequencing to read the CpG methylation and chromatin accessibility across stretches of genomic regions at the single molecule level. Using this approach we have simultaneously determined phased patterns of native methylation and chromatin accessibility in four different human cell lines.

## RESULTS

### Nanopore GpC methylation calling

We previously developed a software tool, nanopolish, which can measure CpG methylation from nanopore sequencing^6^. Specifically, nanopolish employs a hidden Markov Model (HMM) to detect cytosine methylation based on electrical current signatures (events) corresponding to groups of nucleotide sequences (k-mers). The HMM uses a table of event level distributions characteristic to every k-mer, termed a pore model, to predict the methylation state of k-mers. The CpG methylation pore model was generated from sequencing data of DNA enzymatically methylated by M.SssI at >95% of all CpG locations as previously described^6^. The methylation caller outputs log-likelihood ratios for the probability of methylation at a given k-mer, and a threshold is applied to determine the binary value of methylation at single-molecule resolution. To expand the nanopolish algorithm to detect cytosine methylation at GpC contexts, we generated a new training set using combinations of M.SssI (CpG methyltransferase) and M. CviPI (GpC methyltransferase) on unmethylated (PCR amplified) *Escherichia coli (E. coli)* genomic DNA (see Methods). This resulted in samples with CpG, GpC, and both CpG and GpC methylation as well as an unmodified negative control. The methylation samples were sequenced on Oxford Nanopore Technologies (ONT) MinIONs using the sequencing library preparation kit by ligation (LSK-SQK109). The resulting raw data was basecalled using guppy and aligned to the *E. coli* reference genome.

We extracted the event signals and compared the distributions in current for a set of k-mers containing combinations of the methylation motifs. In k-mers that contained motifs for GpC methylation, we observed that the GpC methylated samples had clear shifts in event level distributions in comparison to unmethylated samples (**Fig. 1a**). We further observed that in some k-mers that contain both CpG and GpC motifs, the three methylated samples had different shifts in current, indicating that CpG and GpC methylation could be detected simultaneously in some contexts. The sequencing data from the GpC methylation sample was used to train the pore model for GpC methylation detection.

**Figure 1.**
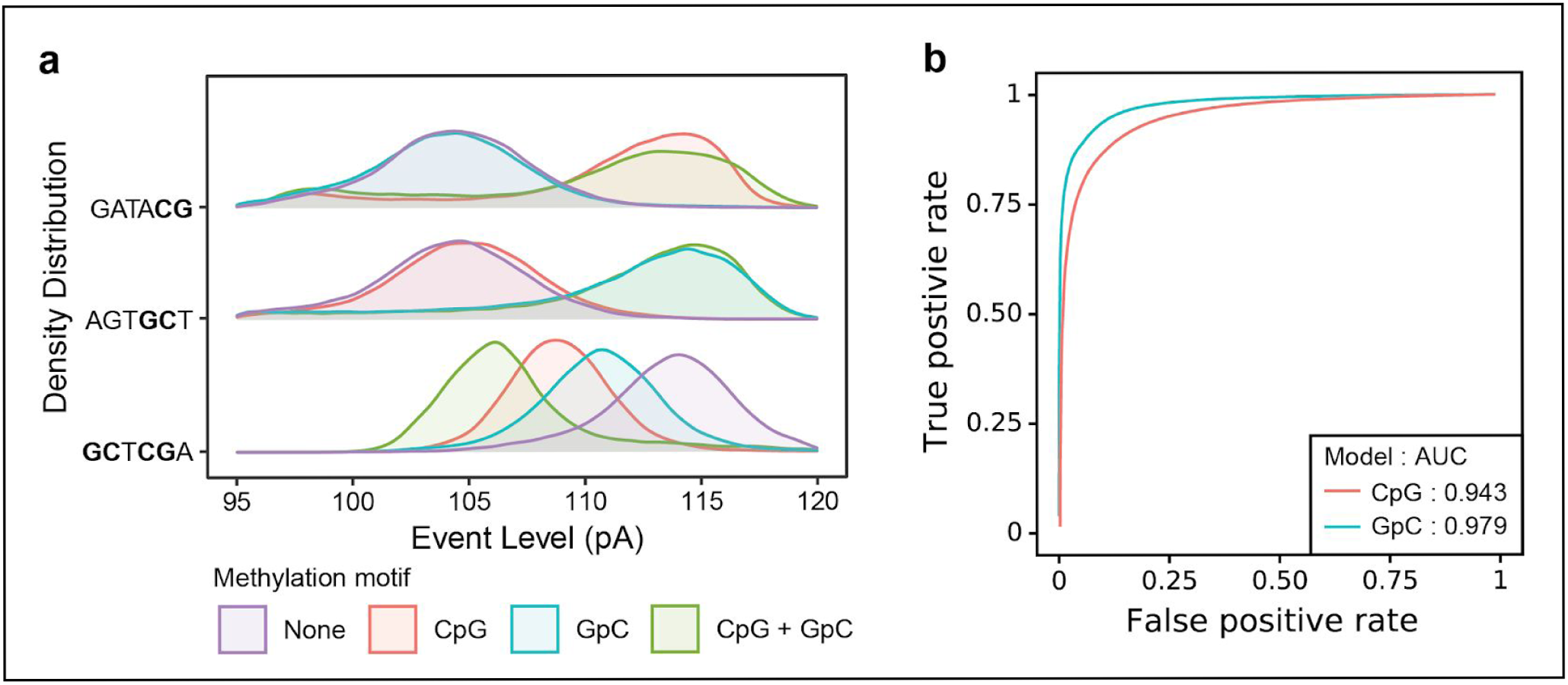
Nanopore sequencing accurately detects GpC methylation. **(a)** Event level current density distribution for select k-mers having CpG, GpC, and both CpG and GpC motif for *E. Coli* genomic DNA with no methylation, CpG, GpC, and both CpG and GpC methylation. **(b)** ROC curve for a range of thresholds for methylation detection on control samples (GM12878 genomic DNA modified with CpG and GpC methylation).

To benchmark GpC methylation detection, we tested methylation pore models on completely methylated and unmethylated samples of genomic DNA from GM12878 human lymphoblast cell line. As with *E. Coli* samples, we generated a completely unmethylated sample of GM12878 gDNA from whole genome PCR-amplification, then treated this with either M. SssI (CpG) or M. CviPI (GpC) to generate the methylated samples. The extent of methylation was validated via low coverage whole genome bisulfite sequencing, using ~2 Mb of sequencing data per condition, which confirmed that the methylated samples were methylated at >95% of all detected motifs (**Supplementary Table 1**). These same validated samples were used as the truth sets for unmethylated, CpG methylated, and GpC methylated events. To assess the performance of methylation detection, we generated receiver operating characteristic (ROC) curves by applying a range of thresholds to bin methylation statuses (**Fig. 1b**) as we did previously^6^. Using a log-likelihood ratio of 2.5 as the threshold for calling methylation (where a value <−2.5 is unmethylated, >2.5 is methylated, and (−2.5, 2.5) is not called), we called 95% of CpGs as methylated in the 72% of all possible CpG k-mers and 97% of GpCs as methylated in 89% of all possible GpC k-mers.(**Supplementary Fig. 1a**). Both the CpG and GpC models had high area under the curve (AUC) of the ROC curve, confirming the applicability of the models. It should again be emphasized that the bisulfite sequencing data indicated incomplete (~96-98%) enzymatic methylation in this sample, so this is a conservative estimate of our accuracy. We used this threshold of 2.5 for subsequent methylation detection.

**Table 1.**
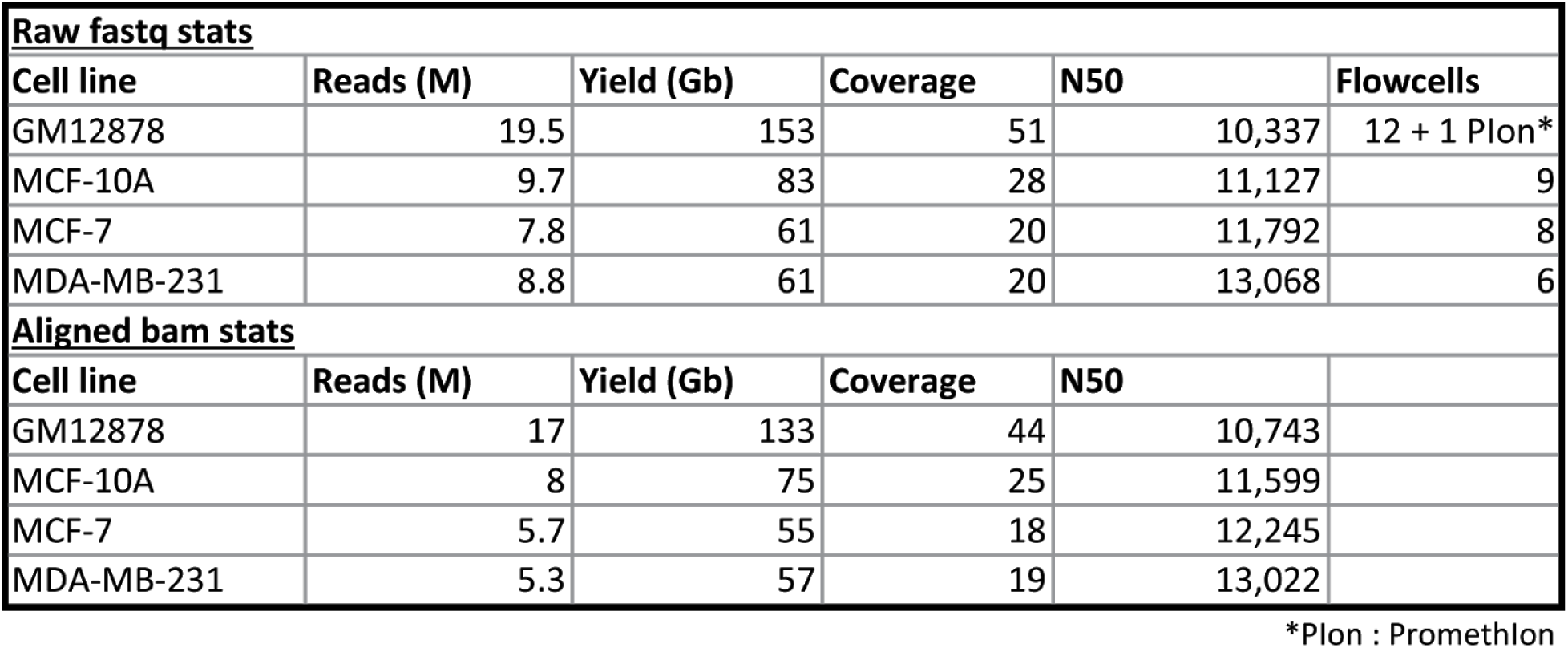
Sequencing statistics of nanoNOMe data. NanoNOMe was performed on four cell lines using multiple runs of MinION, GridION, or PromethION sequencing and pooled to generate one data set per cell line.

### Chromatin and DNA methylation profiling with NanoNOMe

We then adapted the existing NOMe-seq protocol^5^ to profile chromatin state for use with nanopore sequencing, terming this modified method nanoNOMe **(Fig. 2a)**. Because nanopore sequencing discriminates methylated cytosines directly, bisulfite conversion and PCR amplification are unnecessary. However, to preserve the modifications, we cannot amplify the DNA requiring a higher (1-2ug) initial amount of DNA as input. Briefly, intact nuclei were extracted from cells by gentle lysis, followed by methylation with GpC methyltransferase. The methylation treatment of intact nuclei results in GpC methylation only at unoccupied, open regions of the genome (**Fig. 2a**). After purification of DNA from these nuclei by phenol:chloroform extraction and ethanol precipitation, we performed ligation-based library preparation for nanopore sequencing (ONT). After sequencing, basecalling, and alignment, we applied our GpC methylation model to detect GpC methylation in addition to detecting CpG methylation using the existing model. Subsequently, methylation at cytosines in a GCH context were used as a measure of chromatin accessibility and cytosines in a HCG context were used as measures of endogenous methylation, and methylation measurements in GCG cytosines were excluded from analysis. In describing GpCs state, a methylated GpC was interpreted as an accessible mark, and unmethylated as inaccessible.

**Figure 2.**
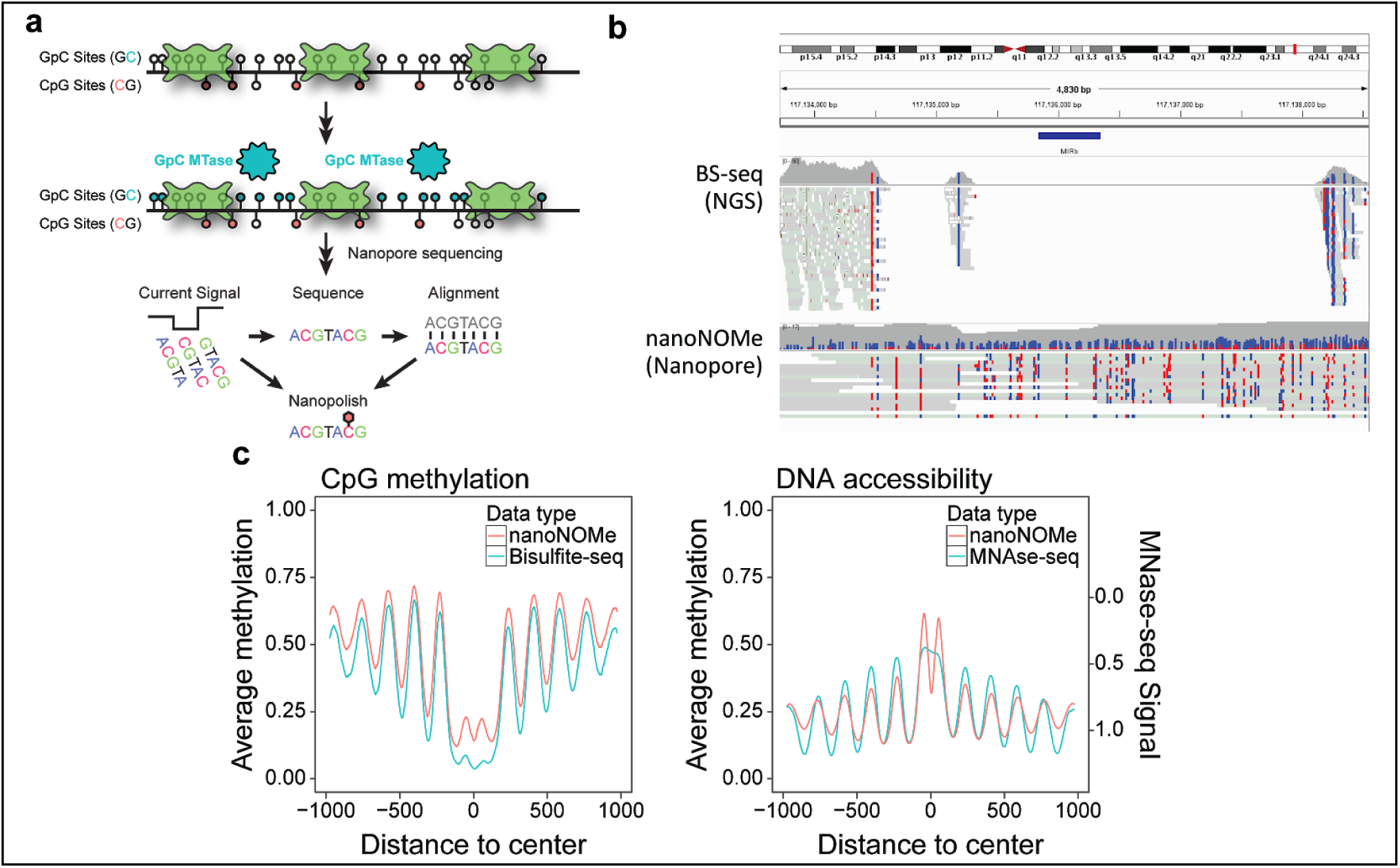
NanoNOMe can measure endogenous DNA methylation and accessibility. (**a**) nanoNOMe uses exogenous methylation of accessible DNA by GpC methyltransferase, followed by nanopore sequencing and nanopolish methylation detection. (**b**) Using bisulfite mode of IGV to visualize methylation on nanopore sequencing reads versus bisulfite sequencing. Note gaps in coverage in highly repetitive region. (**c**) Aggregate plots of endogenous methylation (left) and chromatin accessibility (right) from nanoNOMe (red) and conventional assays (BS-seq and MNase-seq, blue) centered around CTCF binding sites.

We performed nanoNOMe on the GM12878 lymphoblast cell line, chosen because it has been well-characterized in previous studies^8,9^. We generated 128 Gb of mapped sequencing data from 13 flowcells (12 minION and 1 PromethION), with an N50 read length of 10,624 bp. (**Table 1, Supplementary Table 2**). We first compared genomic coverage of the resulting nanoNOMe data to whole genome bisulfite sequencing (WGBS) from a previous study (ENCODE accession ENCSR890UQO). We assessed the ability of nanoNOMe to cover regions that are poorly mappable via short reads by focusing on regions that were enriched in WGBS reads with low mapping score (10 or more reads with mapping quality < 5). These regions covered 132 Mb of the human genome, comprising of 57,982 distinct regions with average size of 2.3 kb. The average coverage of high mapping quality nanoNOMe reads (mapping quality>20) in these regions was 21x, confirming that long read sequencing, and specifically nanoNOMe, is able to cover these regions of low mappability. As an example, we plotted nanoNOMe and WGBS sequencing data in the region surrounding a SINE element, AluSP in chr5:70,903,500-70,904100 (**Fig. 2b**). We observed that the long reads generated from nanopore sequencing provide more even coverage across the region and stretch through the repetitive element, allowing us to measure methylation in and around the entire repetitive element.

We next assessed the performance of nanoNOMe in resolving nucleosome occupancy around CTCF binding sites as done by Kelly, et. al.^5^. We used CTCF binding sites determined by conserved CTCF-binding motifs^10^ that were >2kb away from transcription start sites and experimentally shown to be bound by CTCF in GM12878^9^. We generated aggregate plots of methylation and chromatin accessibility relative to these CTCF binding sites (**Fig. 2c**). The methylation and DNA accessibility agreed with gold standard methods (WGBS and MNase-seq, respectively) from previous studies (**Fig. 2c,** ENCODE accession ENCSR890UQO and ENCSR000CXP). Specifically, we observed that both chromatin accessibility and methylation demonstrate an oscillation in aggregate methylation, propagating from the center of the CTCF binding sites. The distance between the peaks was ~180 bp, corresponding to the typical spacing of mononucleosomes and linker DNA observed near CTCF sites.

### NanoNOMe reveals allele-specific patterns of methylation and nucleosome positioning

We next explored the applicability of long-reads generated from nanoNOMe in detecting patterns of the epigenetic features. Using the epigenetic features encoded on long sequences of reads, we can observe patterns of these features along the length of the reads, e.g. positioning of multiple nucleosomes on single strands of DNA by oscillation of GpC methylation. Using the bisulfite mode on IGV to show GpC methylation (chromatin accessibility), we can view accessibility over the length of long reads at single-read resolution. An exemplar region demonstrating nucleosome occupancy at a CTCF binding site is shown in (**Fig. 3a**). However, the inherent heterogeneity of the chromatin due to the dynamic nature of nucleosome positioning is directly translated to the single-read data, making it difficult to observe patterns of epigenetic features^11^. The biological heterogeneity of DNA accessibility is further compounded by errors associated with the enzymatic methylation, such as imperfect methylation efficacy, non-specific methylation, and dissociation of nucleosomes in a small fraction of DNA during lysis. In order to resolve patterns of methylation and DNA accessibility on single-read resolution, we have to account for the heterogeneity and noise. To that end, we focused on co-occurrences of methylated or unmethylated cytosine on each read, where the co-occurrence is defined by same type of event (methylated or unmethylated) being observed at two distinct positions on a given read (see Methods). Consolidating the co-occurrence across reads in a given region, we found that patterns of read-level nucleosome positioning across the length of reads can be resolved using a co-occurrence matrix (**Fig. 3b**). Because this analysis measures the relationship of (un)methylated cytosine between positions on individual reads, the peaks in the heatmap highlight locations of nucleosome positioning and distances between them, whereas the average plot only indicates that nucleosomes were present at the troughs without any relationship to other troughs along the region.

**Figure 3.**
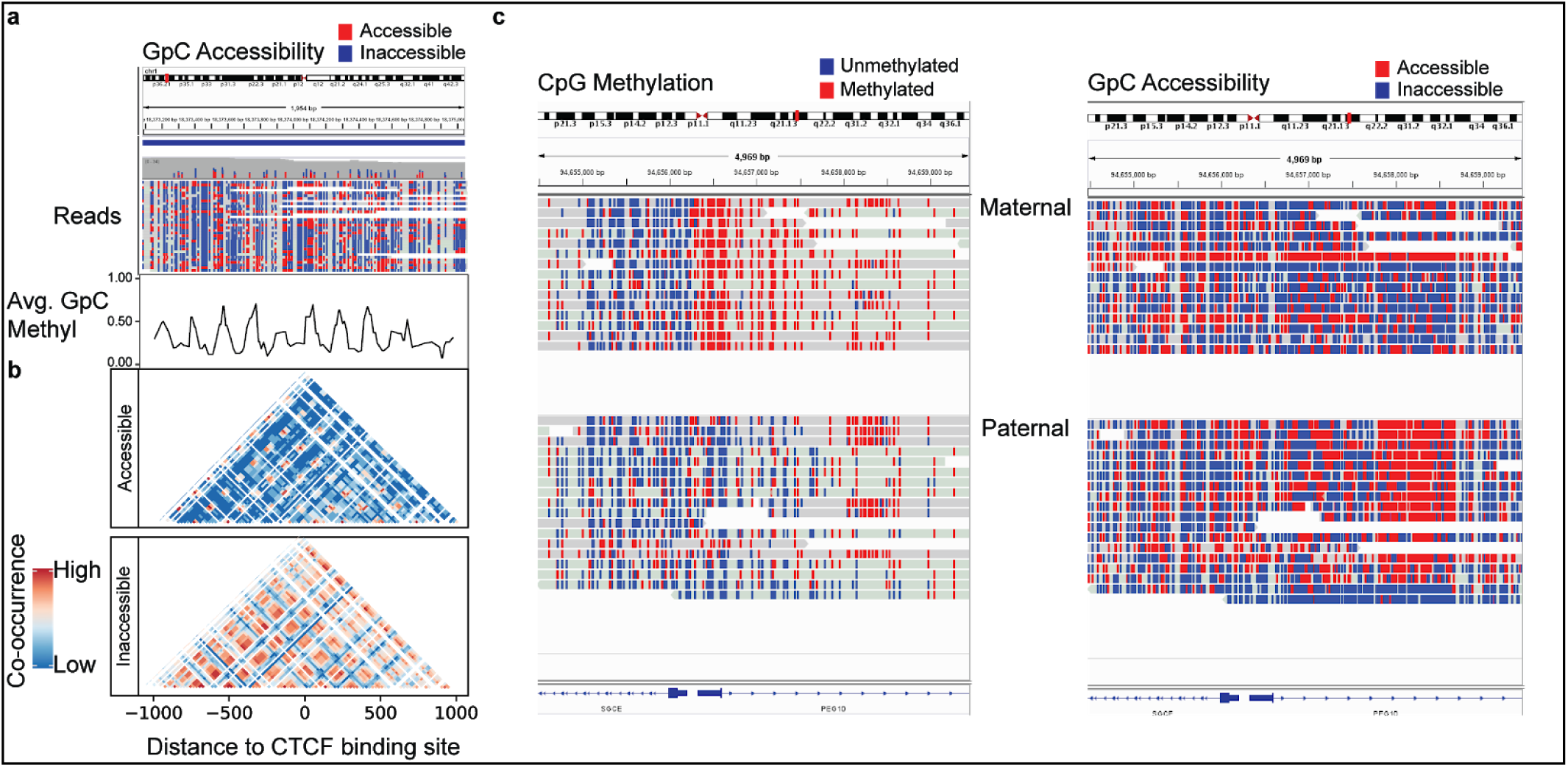
Long single read chromatin and methylation patterns. (**a**) GpC methylation (accessibility) of a CTCF binding site, both average frequency (top) and individual reads (bottom). (**b**) Co-occurrence of accessibility and inaccessibility (GpC) across the lengths of single reads and consolidating co-occurrences across all reads in the region resolves patterns such as nucleosome occupancy. (**c**) Reads mapped to the *PEG10* gene region were separated based on alleles, showing that the reads from paternal, expressed, allele are unmethylated (CpG) at the promoter and open (GpC) in the gene body.

We then used the ability of nanopore sequencing to phase reads into haplotypes^12^ to compare the patterns of chromatin accessibility and methylation between maternal and paternal reads (**Fig. 3c**). Using existing deep coverage data on GM12878 and both parents^13^, we selected heterozygous SNPs belonging to either the maternal or paternal allele. Because nanopore sequencing generates long reads, each read has a greater chance of encountering a heterozygous SNP which can be used to phase the reads into maternal or paternal origin. As an example, we examined the promoter for PEG10 (Paternally expressed gene 10), which is known to be expressed only from the paternal allele^14^. At the PEG10 promoter, we noted a defined region that largely is unmethylated in the paternal allele while being heavily methylated in the maternal allele. We also observe allele specific chromatin accessibility in the promoter-proximal gene body of PEG10, with the paternal allele showing a region of consistently accessible while the maternal allele remains inaccessible (**Fig. 3c**). We used the HOMER (Hypergeometric Optimization of Motif EnRichment) suite of tools^15^ to examine this region of differential accessibility, and revealing that this region is dense in zinc-finger binding motifs, notably the Zn-finger transcription factors KLF5 and KLF14. Members of the KLF family are known to exert both activating and inhibitory activity through chromatin remodeling and recruitment of co-activator or co-repressors^16^, and therefore this region of increased accessibility on the paternal allele may highlight a regulatory element for this imprinted gene PEG10. Interestingly, both of these observations for PEG10 had been predicted by Fang et al ^17^, using computational methods to mine short-read bisulfite sequencing data. Their analysis suggested allele specific methylation at the promoter for PEG10 as well as the presence of intragenic regulatory elements on this gene.

### Epigenome and Gene Expression

We also explored the relationship between epigenetic states and gene expression. Upon generating the metaplots with respect to distance to TSS of annotated genes and stratifying the measurements based on GM12878 gene expression quartiles (ENCODE accession ENCSR843RJV), we observed that chromatin and DNA methylation show shifts in signal depending on expression, where endogenous methylation decreases at promoter regions with increasing expression and accessibility increases with increasing expression (**Supplementary Fig. 2**). To observe the epigenetic states for each gene and measure how the two features are directly related with respect to gene promoters, we calculated the average endogenous methylation and accessibility in individual promoter regions (400 bp window around transcription start sites) (**Fig. 4a**). We observed that promoters of genes with low expression tend to be highly methylated with low accessibility, and with increasing expression, the cluster shifted first to lower methylation and low accessibility, then to low methylation and higher accessibility. These results suggest that a combination of accessibility and methylation may be more useful to understand gene regulation than either independently.

**Figure 4.**
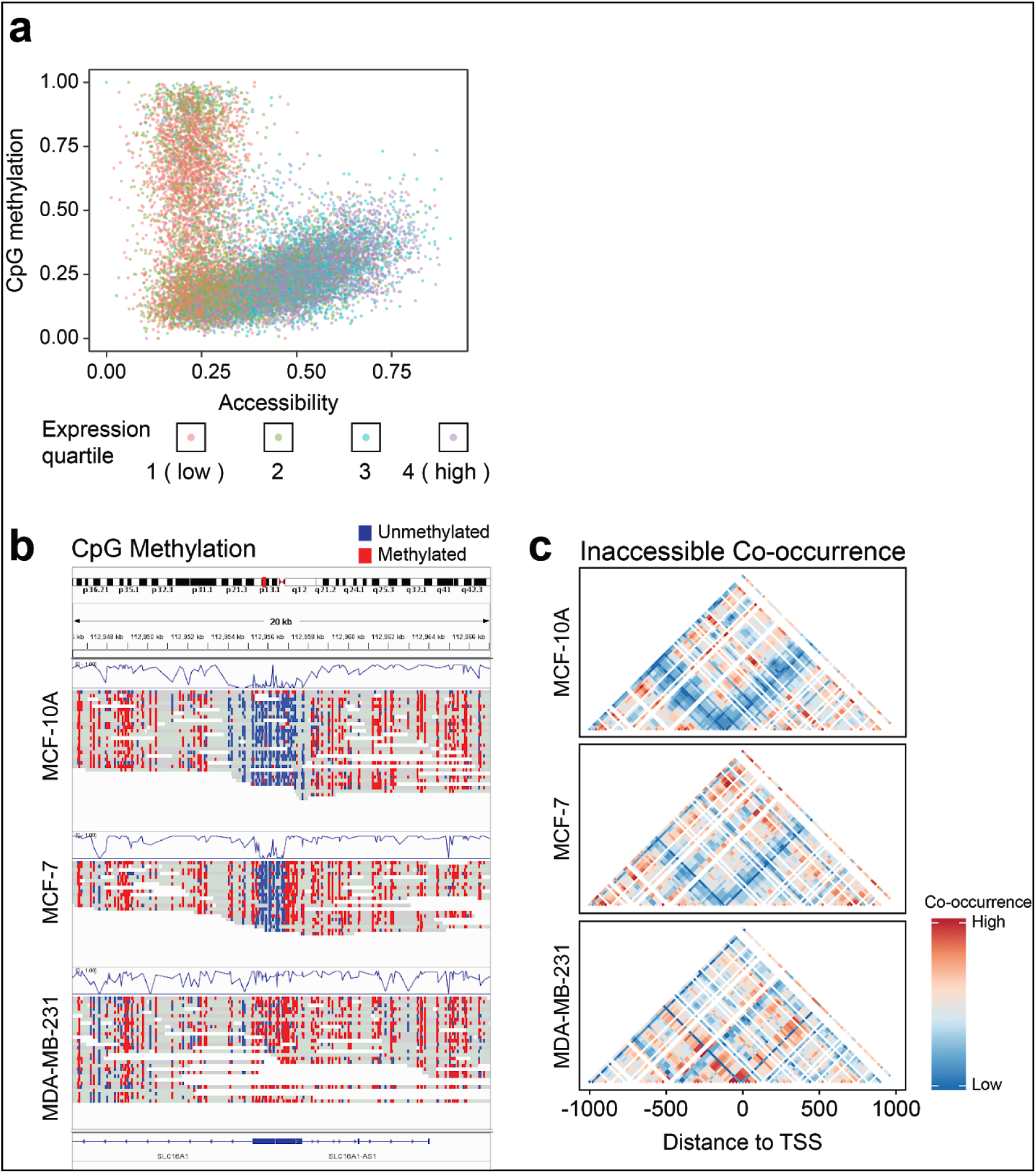
Epigenetic state correlation with gene expression. (**a**) Scatter plots of endogenous methylation versus accessibility color-coded by expression quartile in GM12878. Each dot in the scatter plot represents the methylation and chromatin state of a promoter region of a gene (200 bp +/- TSS) (**c**) IGV view of endogenous methylation surrounding *SLC16A1* shows the differential methylation around the promoter region of the gene and (**d**) co-occurrence plot of accessibility and inaccessibility showing increased co-occurrence of chromatin inaccessibility around the promoter.

Next, we explored the applicability of nanoNOMe on differential epigenetics analysis by performing nanoNOMe on three well-characterized breast cell lines: MCF-7 (luminal breast carcinoma, ER+/PR+/HER2-) and MDA-MB-231 (basal breast carcinoma, ER-/PR-/HER2-) as two subtypes of breast cancer, and MCF-10A (fibrocystic disease) as the normal baseline subtype (**Table 1**)^18,19^. We achieved >20x whole genome coverage of nanoNOMe data per cell line. Comparison of epigenetic states on promoter regions of differentially expressed genes revealed that a decrease in endogenous methylation coupled with an increase in accessibility is reflective of an increase in transcription; while an increase in methylation coupled with a decrease in accessibility is correlated with a decrease in transcription (**Supplementary Fig. 3**). In all three comparisons, higher methylation and lower accessibility favors decrease in expression (under-expression), and lower methylation and higher accessibility favors increase in expression (over-expression).

We then examined SLC16A1, one of the genes down-regulated in the cancer cell lines, that also exhibited differential methylation and chromatin state(**Fig 4b,c**). The methylation frequency confirms that the promoter region of SLC16A1 is largely unmethylated in MCF-10A and methylated in MCF-7 and MDA-MB-231. The single-read data accurately captures the cell-to-cell variability in methylation, where a few of the reads in MCF-10A exhibit methylation across the entire promoter region even when the vast majority of reads are unmethylated. We can clearly observe the erosion of the 3kb unmethylated region from the normal (MCF10A) to the cancer (MCF-7) to the aggressive cancer (MDA-MB-231) cell line, all on individual reads. Turning to chromatin accessibility, MCF-10A had a relatively wide region of accessible chromatin around the TSS whereas MCF-7 had a narrow window and MDA-MB-231 is completely inaccessible. Furthermore, the read-level co-occurrence revealed that MCF-7 had a strong frequency of co-occurrence of inaccessibility −1000 bp upstream and downstream of the TSS, suggesting blocking of transcription by occupancy up and downstream of the TSS, such as chromatin looping, whereas in MDA-MB-231, the down-regulation occurred by total occupancy of the TSS (**Fig. 4d**)

### Structural Variations and the Epigenome

We also detected structural variations and compared nanoNOMe patterns at these sites across the three breast cell lines(**Supplementary Table 2**)^20,21^. We called a total of 25,882 SVs across all three breast lines and compared these using SURVIVOR^21^. The most abundant variant type were deletions (13,974) followed by insertions (10,127). The majority of the SVs were singletons (53.8%) with 28.3% overlapping over two samples and 17.8% all three samples. The majority of genes (67.36%) impacted by SVs were from deletions.

Selecting just SVs that occur only in the cancer cell lines (MCF-7 and MDA-MB-231) but not in the normal breast cell line (MCF-10A), we examined the epigenetic state of the regions flanking breakpoints. We found that regions flanking the breakpoints of structural variations do not exhibit consistent epigenetic characteristics in these cell lines, suggesting that structural variations have complex epigenetic consequences, dependent on more factors than the type of the variant (**Supplementary Fig. 3**). For example, in a 4,500 bp homozygous deletion in chr19 of MCF-7 both flanking regions are strongly hypermethylated as compared to the normal cell lines while MDA-MB-231 has milder hypermethylation. (**Fig. 5a**). The read-level co-occurrence chromatin occupancy in MCF-7 showed that the regions around the deletion are less frequently coordinated but instead oscillate in occupancy, indicating protection of the deletion by positioning of nucleosomes around the breakpoints. In MDA-MB-231 the entire flanking region is marked by coordinated inaccessibility (**Fig. 5b**).

**Figure 5.**
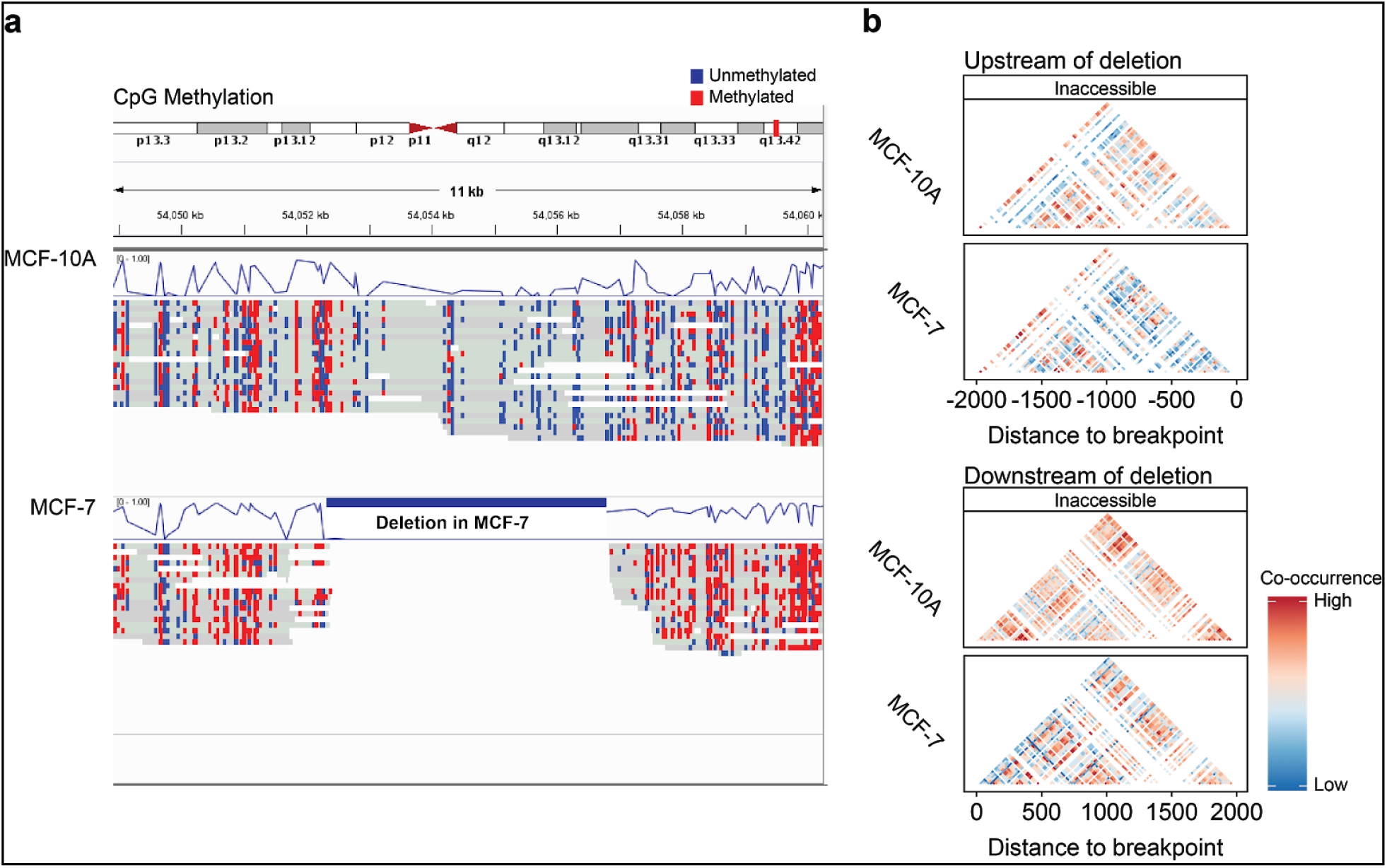
Epigenetic state comparison of homozygous deletion present only in MCF-7 and MDA-MB-231. (**a**) A deletion in IGV view shows that the flanking regions around the deletion in MCF-7 are hypermethylated compared to MCF-10A while MDA-MB-231 is not, and (**b**) co-occurrence matrix of inaccessible events indicates MCF-7 has less consistent inaccessibility (more blue) and more nucleosome occupancy around the breakpoints (oscillation in color) whereas MDA-MB-231 is marked by increased inaccessibility up and downstream.

## DISCUSSION

We have demonstrated a method to examine endogenous methylation and chromatin accessibility on long fragments of DNA. Leveraging long reads we can measure allele specific methylation and chromatin profiles. We have also shown that because nanopore sequencing reads span multiple nucleosomes, nucleosome occupancy on single reads can be observed and the frequency of nucleosome positioning can be determined to observe consistency and patterns of nucleosome positioning. Using existing expression data for GM12878, we evaluated how chromosome accessibility and cytosine methylation were related to gene transcription. We found that these features were able to explain part of the differences in gene transcription, with the overall trend of the most highly expressed genes demonstrated the most accessible promoters with the lowest amount of CpG methylation. Extending our studies into breast cell lines, we compared the epigenetic state of the promoters of differentially expressed genes and found that focusing on read-level data allowed us to discern patterns of nucleosome occupancy. Further, we can observe epigenetic states of alleles with structural variants, allowing combined measurement of large genetic mutation and epigenetic state with the same assay.

However, this method is still limited, in part by using the same methylation mark that already exists in mammalian cells, 5-methylcytosine. Moving forward, we can take advantage of other methyltransferases, e.g. EcoGII which methylates adenine to N6-methyladenine. Such a technique could also provide a “multi-color” measurement, allowing further aspects of the epigenome to be integrated on the same molecule. Others have already leveraged this methyltransferase fused to lamin protein^22^, to explore nuclear architecture but are limited to immunoprecipitation based sequencing, precluding single molecule resolution. With further training and development, we may be able to leverage exogenous labeling with nanopore sequencing to store information about the cell state on the DNA, then sequencing it to gain long-range, phased information.

## METHODS

### GpC methylation model generation for nanopolish

Along with the GpC methylation model, the CpG methylation model was also regenerated to ensure the validity of the method for model generation. Genomic DNA from *E. coli* K12 MG1655 (ATCC 700926DQ) and genomic DNA from GM12878 lymphoblast cell line (Coriell Institute) were first sheared to an average fragment size of 8 kb using g-tubes (Covaris Cat. 520079). The fragmented DNA was PCR amplified to generate unmethylated DNA using the first steps of low input ligation kit SQK-LWP001 (ONT). Briefly, samples were end-repaired, deoxyadenosine(dA)-tailed, and ligated to amplification adaptors, followed by 11 cycles of PCR amplification. The resulting unmethylated, sheared DNA was methylated with M. SssI (NEB Cat. M0226) for CpG methylation or M. CviPI (NEB Cat. M0227) for GpC methylation, or both enzymes for CpG+GpC methylation. Two cycles of 4-hour methylation were performed for each sample, and for each cycle of treatment S-adenosylmethionine (SAM) and the enzyme were replenished at the 2 hour mark to maximize methylation levels.

### Validation of DNA methylation by bisulfite sequencing

Near-complete methylation in the training samples (*E. coli*) and testing samples (GM12878) were validated by performing whole genome bisulfite sequencing on the Illumina MiSeq platform. NEBnext Ultra library preparation kit (NEB Cat. E7370) and Zymo EZ DNA methylation-lightning kit (Zymo Cat. D5030) were used to generate the bisulfite sequencing libraries. Briefly, DNA from each sample was shared to 300 bp fragments using Bioruptor Pico (Diagenode), followed by end-repair and dA-tailing. Methylated universal adaptor (NEB Cat. E7535) was ligated using the Blunt/TA ligase from the kit. The adaptor-ligated samples were bisulfite-converted, quenched, and cleaned-up before PCR amplification with multiplexing primers and uracil-tolerant Taq polymerase (KAPA HiFi Uracil+ (Roche Cat. KK2801)). The resulting DNA sequencing library was sequenced on an Illumina MiSeq device using a V2 300-cycle chemistry.

The resulting data was analyzed using Bismark version 0.19.0^23^. After alignment, PCR duplicates were removed using Picard tools MarkDuplicates module (http://broadinstitute.github.io/picard/). Reads were truncated at the 3’ end to a max length of 50 bp to minimize any methylation bias at 3’ ends of reads associated with the low complexity bisulfite converted libraries. The total number of methylated cytosine residues and unmethylated cytosine residues were counted to calculate methylation percentages of the samples.

### Cell culture

GM12878 lymphoblast cells were obtained from Coriell Institute and MCF-10A, MCF-7, and MDA-MB-231 breast cells were obtained from ATCC. GM12878 were grown in RPMI 1640 medium (Gibco Cat. 11875119) supplemented with 15% fetal bovine serum (FBS, Gibco Cat. 26140079) and 1% penicillin streptomycin (P/S, Gibco Cat. 15140122). MCF-10A were grown in in DMEM F-12 medium (Gibco Cat. 11320033) supplemented with 5% horse serum (Gibco Cat. 16050122), 10 μg/mL human insulin (Sigma Aldrich Cat. 19278), 20 ng/mL hEGF (Gibco Cat. PHG0311L), 100 ng/mL Cholera toxin (Sigma Aldrich Cat. C8052), 0.5 μg/mL Hydrocortisone (Sigma Aldrich Cat. H0135), and 1% P/S. MCF-7 and MDA-MB-231 were grown in DMEM (Gibco Cat. 11965118) supplemented with 10% FBS and 1% P/S.

### Nucleosome footprinting via GpC methyltransferase

NOMe-seq was performed to the cells with adjustments for nanopore sequencing. Cells were collected by trypsinization, then nuclei were extracted by incubating in resuspension buffer (100 mM Tris-Cl, pH 7.4, 100 mM NaCl, 30 mM MgCl_2_) with 0.25 % NP-40 for 5 minutes on ice. Intact nuclei were collected by centrifugation for 5 minutes at 500xg at 4 °C. Nuclei were subjected to a methylation labeling reaction using a solution of of 1x M. CviPI Reaction Buffer (NEB), 300 mM sucrose, 96 μM S-adenosylmethionine (SAM; New England Biolabs, NEB), and 200 U M. CviPI (NEB) in 500 μL volume per 500,000 nuclei. The reaction mixture was incubated in 37 °C with shaking on thermomixer at 1,000 rpm for 15 minutes. SAM was replenished at 96 μM at 7.5 minutes into the reaction. The reaction was stopped by addition of equal volume of stop solution (20 mM Tris-Cl, pH 7.9, 600 mM NaCl, 1% SDS, 10 mM disodium EDTA). Samples were treated with proteinase K (NEB) at 55 °C for > 2 hours, and DNA was extracted via pheol:chloroform extraction and ethanol precipitation. After proteinase K treatment, and in all following steps, samples were handled with care using large orifice pipette tips to avoid excessive fragmentation of DNA.

### Nanopore sequencing

Purified gDNA was prepared for nanopore sequencing following the protocol in the genomic sequencing by ligation kit LSK-SQK108 (ONT). Samples were first sheared to ~10 kb using G-tubes (Covaris): by centrifuging 2-3 μg of unfragmented gDNA at 5,000x g for 1 minute, then inverting the tube and centrifuging again. We sheared the DNA to 10 kb because it produces long fragments of DNA while maximizing the yield of nanopore sequencing. Shearing to larger sizes or unsheared DNA may be used to maximize the length of sequenced reads, with the caveat that sequencing yield will drop. The sheared samples were end-repaired and dA-tailed using NEBnext Ultra II end-repair module (NEB), followed by clean-up using 1x v/v AMPure XP beads (Beckman Coulter). Sequencing adaptors, comprised of leader adaptor DNA and motor proteins, were ligated to the end-prepared DNA fragments using Blunt/TA Ligase Master Mix (NEB), followed by clean-up using 0.4x v/v AMPure XP beads and sequencing kit reagents. >400 ng of adaptor ligated samples per flow cell were loaded onto FLO-MIN106 or PRO-002 flowcells and run o n MinION Mk1b, GridION, or PromethION sequencers for up to 72 hours.

### Training nanopolish methylation calling for GpC methylation

The nanopore sequencing data from GpC methylated *E. coli* gDNA was used to generate the methylation model with the newest version of nanopolish (v. 0.11.0). This version has updated calculations to improve numerical stability for large datasets. We used the unmethylated and GpC methylated GM12878 gDNA as truth sets to generate the ROC curves and validate the model.

### Data preprocessing (basecalling, alignment, and methylation calling)

Raw current signals were converted to DNA sequences using guppy version 2.1.3 (ONT), using basecalling configuration designed to reduce false-positive deletions in the resulting sequences ^24^. DNA sequences were aligned to hg38 human reference genome without alternative contigs using NGM-LR^20^. CpG and GpC methylation were called using nanopolish version 0.11.0 using a log-likelihood threshold of 2.5 to determine methylation states. We used Sniffles^20^ with default parameters to infer SVs across each sample and SURVIVOR^21^ merge to obtain a multi sample VCF file.

### Comparison of nanoNOMe with conventional methodologies

Bisulfite sequencing data of GM12878 was obtained from ENCSR890UQO, and was processed using Bismark version 0.19.0. After alignment to hg38 reference genome, duplicate reads were removed using Picard tools MarkDuplicates module (http://broadinstitute.github.io/picard/) before further bismark processing to yield methylation frequency values. Normalized MNase-seq signals were obtained from ENCSR000CXP. Methylation frequency and normalized MNase-seq signal at regions surrounding genomic features of interest were extracted for the generation of the aggregate plots. For each genomic feature, average methylation frequency and accessibility was calculated by aggregating methylation calls with respect to distance from the feature and taking the rolling average with a window of 50 bp. Known TSS and CGI were obtained from Gencode (release v29). CTCF binding sites were determined by overlapping computationally predicted CTCF binding sites^10^ with conservative IDR peaks in ChIP-seq of CTCF on GM12878 (ENCODE accession ENCSR000AKB) and removing peaks that fell within 2kb of known TSS.

### Read-level methylation visualization

The bisulfite view setting of IGV was used to visually compare methylation states between samples. In order for this software to function properly, we converted all non-methylated cytosines to thymines (as would occur during bisulfite conversion). All cytosines that nanopolish called as methylated (in either CpG or GpC context) were kept as cytosines. This permitted inspection of endogenous methylation as well as chromatin accessibility using IGV.

To observe patterns of DNA methylation and accessibility in the presence of biological heterogeneity and technical variability, co-occurrence of methylated/unmethylated cytosine is calculated across reads that map to the genomic region of interest. Co-occurrence, c, is defined by the same event, M (methylated or unmethylated), occurring on two separate binned locations, i and j, along a given read:

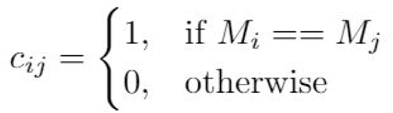

After calculating the co-occurrence for each pair of coordinates for each read, the counts are piled up to determine the frequency of co-occurrence as a measure of how often reads have the same events occurring between the positions i and j. The resulting matrix of co-occurrence pileup is normalized by the maximum count, and plotted as a 2-dimensional heatmap to visualize the patterns (see accessions for code availability).

### Haplotype Assignment and Allele-Specific Methylation Analysis

We obtained genotype information for GM12878 from existing phased Illumina platinum genome data generated by deep sequencing of the cell donors’ familial trio^13^. The bcftools package was used to filter for only variants that are heterozygous in GM12878. We then used the SnpEff^25^ variant annotation and effect prediction tool was used to associate gene names to each read. Starting with aligned reads, we used the extractHAIRS utility of the haplotype-sensitive assembler HapCUT2^26^ to identify reads with allele-informative variants. For allelic assignment, we required a read to contain at least two variants, and required that greater than 75% of identified variants agreed on the parental allele of origin -- this stringent threshold was selected to reduce the chances of incorrect assignment from nanopore sequencing errors. Through this approach each read was annotated as maternal, paternal or unassigned.

## Accessions

NanoNOMe data of GM12878, MCF-10A, MCF-7, and MDA-MB-231 are available at NCBI Bioproject ID PRJNA510783 (http://www.ncbi.nlm.nih.gov/bioproject/510783). Source code is available at https://github.com/timplab/nanoNOMe.

## Acknowledgements

We thank K. D. Hansen for helpful discussions. This study was supported by National Human Genome Research Institute (NHGRI project 5R01HG009190).

