## Supplemental Materials for "Simultaneous profiling of chromatin accessibility and methylation on human cell lines with nanopore sequencing"

SUPPLEMENTAL MATERIAL

### TABLE OF CONTENTS

#### Figures

**Supplementary Figure 1.** Performance and comparison of methylation pore models

**Supplementary Figure 2.** NanoNOMe data in TSS stratified by expression quartile.

**Supplementary Figure 3.** Endogenous methylation and chromatin accessibility in promoter regions.

**Supplementary Figure 4.** Scatterplots of epigenetic comparison around structural variations.

#### Tables

**Supplementary Table 1.** CpG and GpC methylation rates of samples treated with combinations of methylations.

**Supplementary Table 2.** Individual nanopore sequencing run statistics

**Supplementary Table 3.** Summary of structural variations detected in breast cell lines.

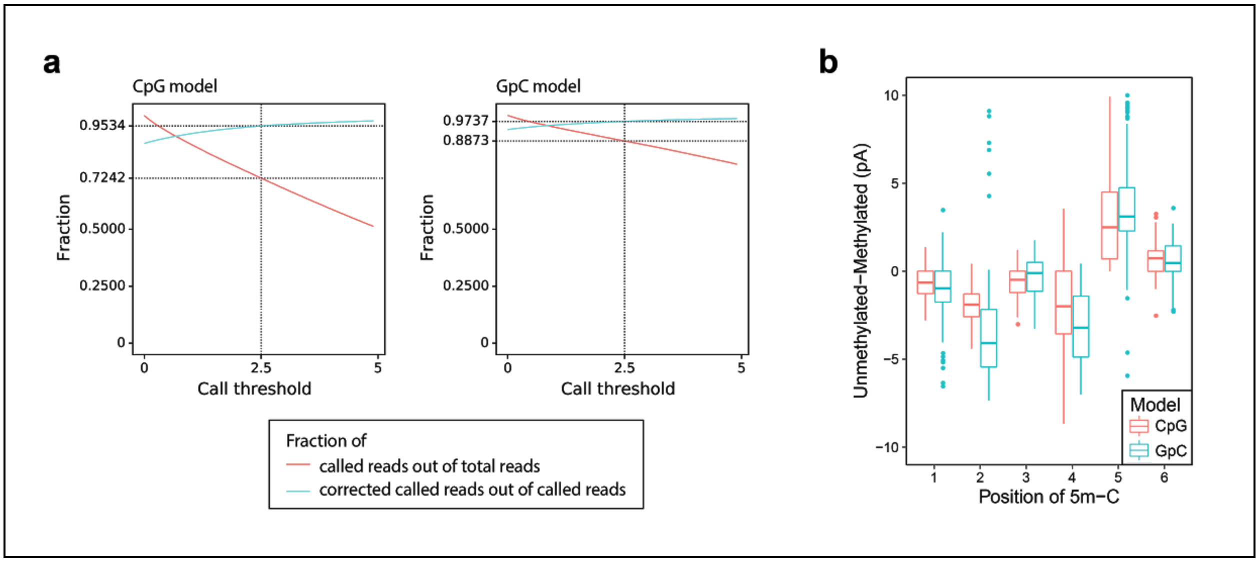


**Supplementary Figure 1. Performance and comparison of methylation pore models.** Performance of CpG and GpC methylation models were measured by (a) measuring the fraction of k-mers passing the threshold filter and the fraction of k-mers from which methylation was correctly called. (b) Influence of position of methylation along the k-mer is determined by calculating the difference of event level mean between methylated k-mers their unmethylated counterparts, and grouping the differences based on the methylation motif position.

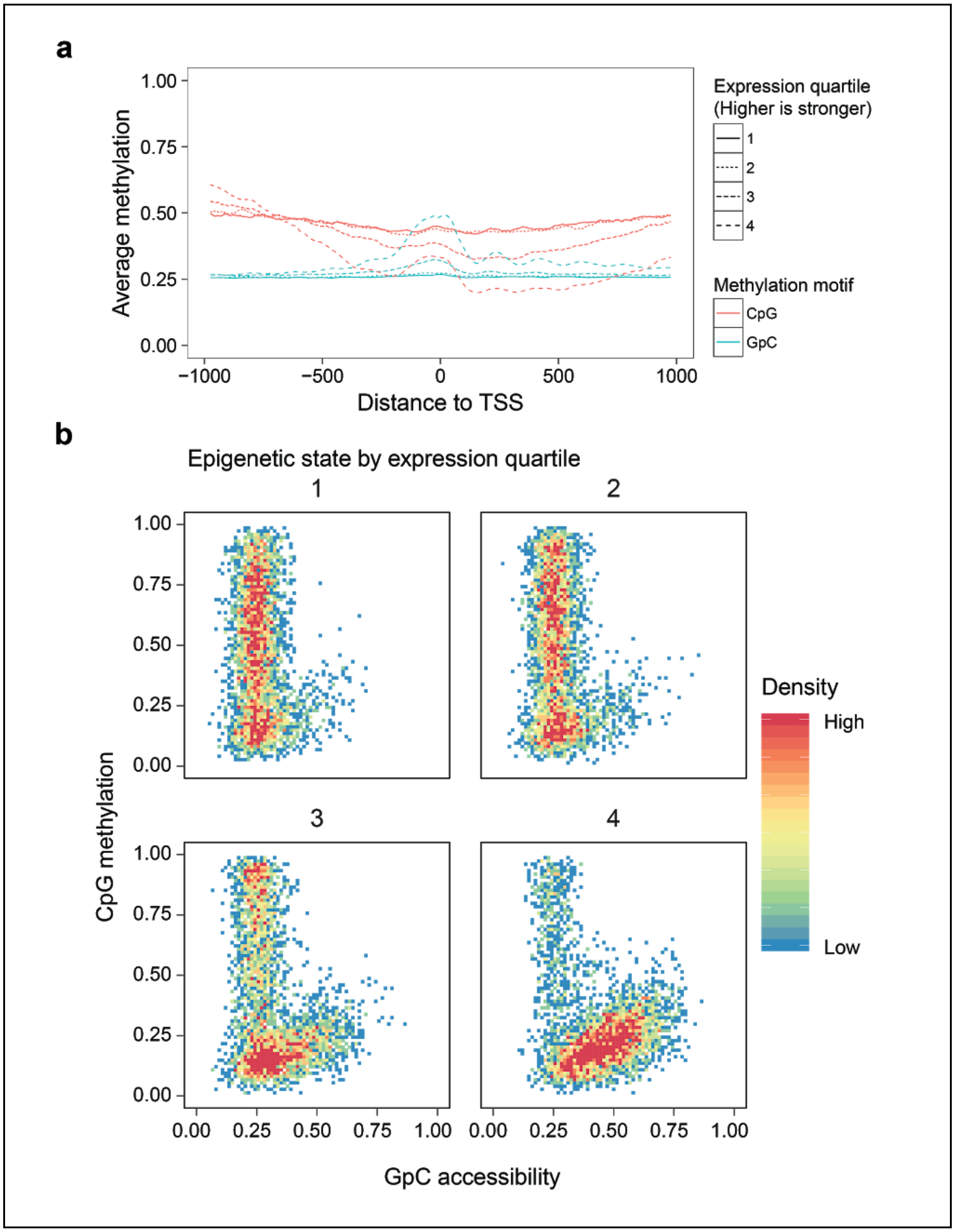


**Supplementary Figure 2. NanoNOMe data in TSS stratified by expression quartile.** (**a**) Metaplot of average methylation and accessibility as a function of distance to transcription start sites, divided up into expression quartiles. (**b**) 2D density plots of the scatter plot ( **Fig 2d**) with each panel representing an expression quartile.

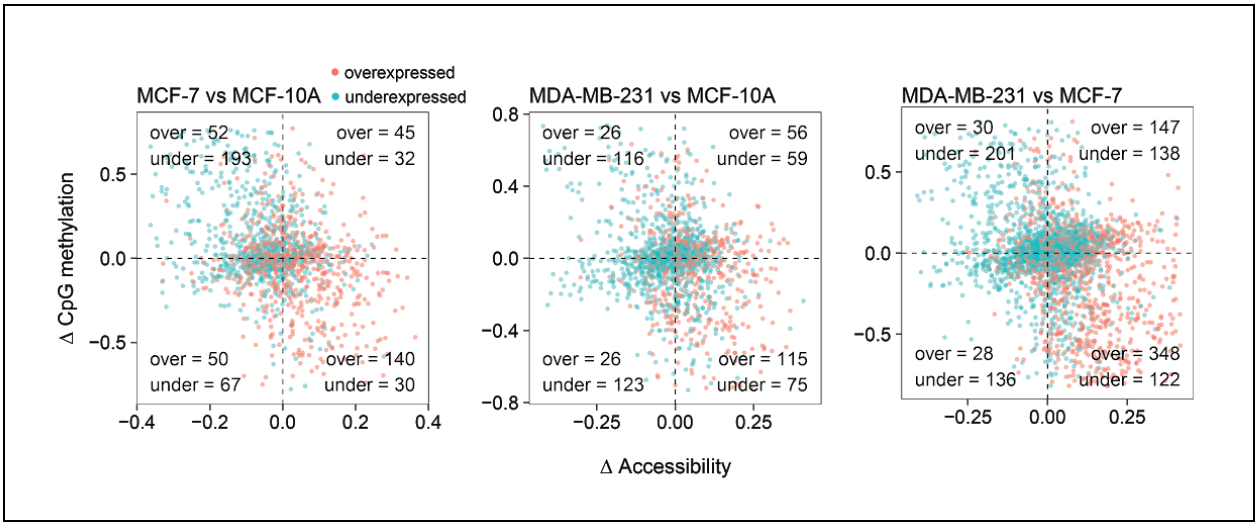


**Supplemental Figure 3: Endogenous methylation and chromatin accessibility in promoter regions** Comparing differentially expressed genes are across three cell lines (MCF-7-MCF-10A, MDA-MB-231-MCF-10A, MDA-MB-231-MCF-7) with genes higher expressed in cancer lines as red and lower expressed in cancer lines as blue. X-axis is chromatin accessibility and Y-axis DNA CpG methylation in the +/- 200 bp window around the TSS of the gene. The numbers in each quadrant are the number of over-expressed genes and under-expressed genes in the given quadrant.

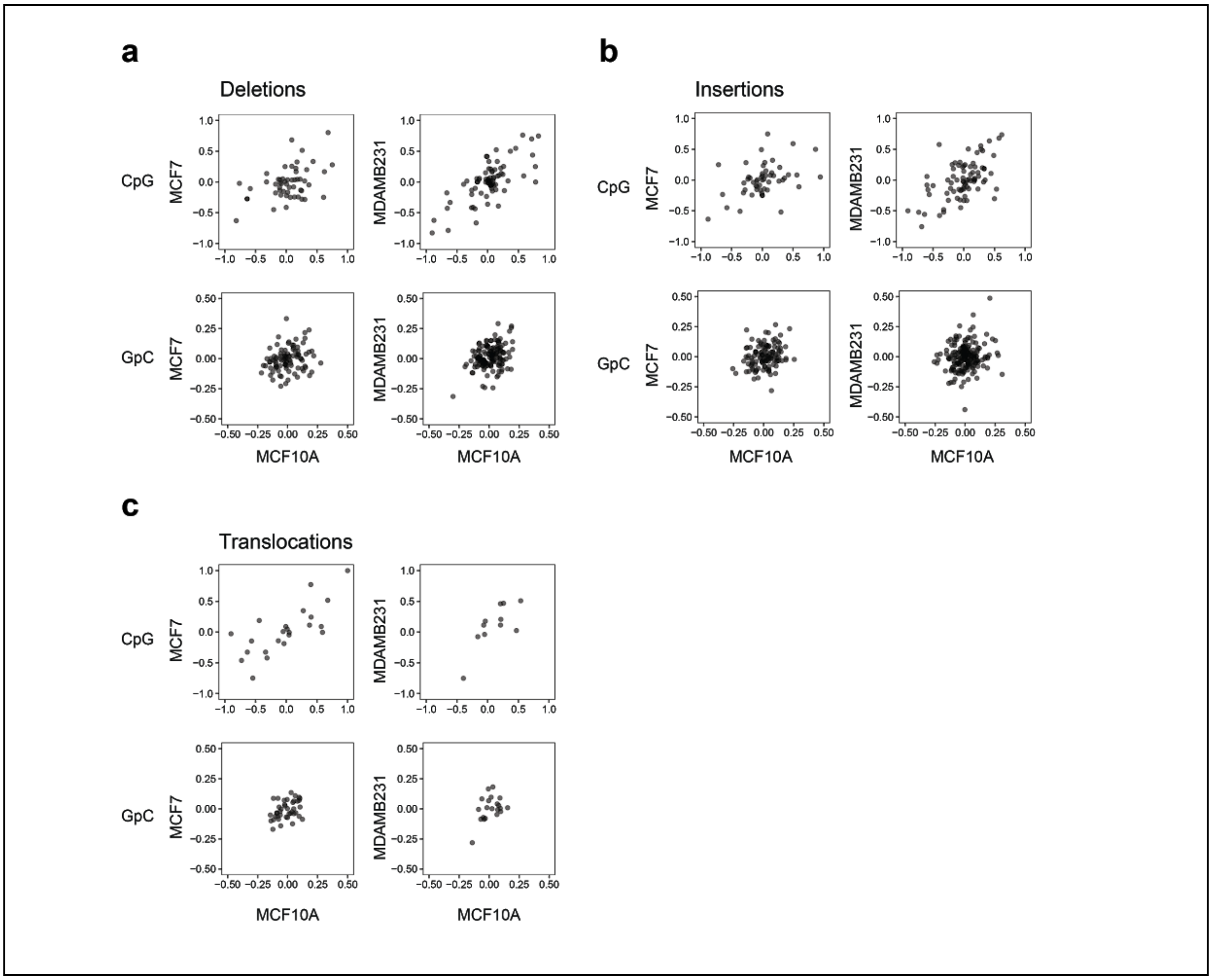


**Supplementary Figure 4. Scatterplots of epigenetic comparison around structural variations.** Scatterplots of difference in methylation (top panels) and accessibility (bottom panels) between 5' and 3' flanking regions of breakpoints (BP2-BP1) of different types of structural variations ((**a**) deletions, (**b**) insertions, (**c**) translocations) that occur in cancer cell lines (MCF-7 and MDA-MB-231, y-axes) but not in MCF-10A (x axes).

| Origin | Methylation | Detected methylation | Methylated loci | Unmethylated loci | Methylation % |
|---|---|---|---|---|---|
| Ecoli | CpG | CpG | 278724 | 5446 | **98** |
| Ecoli | CpG | GpC | 10629 | 485977 | **2** |
| Ecoli | CpGGpC | CpG | 235883 | 4051 | **98** |
| Ecoli | CpGGpC | GpC | 430714 | 7089 | **98** |
| Ecoli | GpC | CpG | 12080 | 172313 | **7** |
| Ecoli | GpC | GpC | 359733 | 5412 | **99** |
| Ecoli | Unmethylated | CpG | 4132 | 158333 | **3** |
| Ecoli | Unmethylated | GpC | 3529 | 292160 | **1** |
| GM12878 | CpG | CpG | 210033 | 8703 | **96** |
| GM12878 | CpG | GpC | 19158 | 1866755 | **1** |
| GM12878 | CpGGpC | CpG | 179562 | 8364 | **96** |
| GM12878 | CpGGpC | GpC | 1628962 | 42718 | **97** |
| GM12878 | GpC | CpG | 10838 | 243231 | **4** |
| GM12878 | GpC | GpC | 2374964 | 48817 | **98** |
| GM12878 | Unmethylated | CpG | 2478 | 195918 | **1** |
| GM12878 | Unmethylated | GpC | 13620 | 1739225 | **1** |

**Supplementary Table 1. CpG and GpC methylation rates of samples treated with combinations of methylations.** Total number of methylated loci and unmethylated loci for each sample was tabulated to calculate the percent methylation per sample.

| Cell | Experiment number | Flowcell type | Number of flowcells | Number of raw reads (M) | Total raw bases (Gb) | Aligned reads (M) | Aligned bases (Gb) | N50 length |
|---|---|---|---|---|---|---|---|---|
| GM12878 | 1 | FLO-MIN106 | 2 | 1.64 | 12.95 | 1.30 | 11.20 | 11,475 |
| GM12878 | 2 | FLO-MIN106 | 2 | 1.90 | 14.86 | 1.56 | 12.68 | 10,736 |
| GM12878 | 3 | FLO-MIN106 | 2 | 0.81 | 6.53 | 0.68 | 5.66 | 10,526 |
| GM12878 | 4 | FLO-MIN106 | 2 | 1.03 | 11.10 | 0.89 | 9.79 | 17,346 |
| GM12878 | 5 | FLO-MIN106 | 2 | 1.95 | 14.77 | 1.55 | 12.11 | 11,635 |
| GM12878 | 6 | FLO-MIN106 | 2 | 2.09 | 15.42 | 1.69 | 13.55 | 11,156 |
| GM12878 | 7 | FLO-PROM002 | 1 | 11.51 | 72.84 | 8.90 | 62.62 | 9,791 |
| MCF10A | 1 | FLO-MIN106 | 2 | 0.86 | 6.52 | 0.60 | 5.70 | 11,428 |
| MCF10A | 2 | FLO-MIN106 | 3 | 3.59 | 33.11 | 3.05 | 29.64 | 11,888 |
| MCF10A | 3 | FLO-MIN106 | 4 | 4.96 | 41.96 | 4.09 | 37.06 | 11,215 |
| MCF7 | 1 | FLO-MIN106 | 1 | 1.17 | 9.71 | 1.03 | 8.77 | 11,210 |
| MCF7 | 2 | FLO-MIN106 | 2 | 0.78 | 5.29 | 0.52 | 4.72 | 11,652 |
| MCF7 | 3 | FLO-MIN106 | 5 | 5.42 | 44.25 | 4.48 | 39.61 | 12,462 |
| MCF7 | 4 | FLO-MIN106 | 3 | 1.62 | 17.51 | 1.46 | 16.05 | 18,532 |
| MDAMB231 | 1 | FLO-MIN106 | 1 | 0.89 | 8.22 | 0.78 | 7.39 | 11,531 |
| MDAMB231 | 2 | FLO-MIN106 | 2 | 2.10 | 19.01 | 1.82 | 17.30 | 11,488 |
| MDAMB231 | 3 | FLO-MIN106 | 3 | 3.09 | 33.06 | 2.68 | 30.14 | 14,583 |
| MDAMB231 | 4 | FLO-MIN106 | 3 | 1.87 | 22.10 | 1.67 | 20.06 | 15,783 |

**Supplementary Table 2. Individual nanopore sequencing run statistics.**
Nanopore sequencing run statistics for individual sequencing experiments performed, before pooling them by cell line to achieve the final yields

| Cells conatining SV | DEL | TRA | DUP | INV | INS | Total |
|---|---|---|---|---|---|---|
| Only MCF-10A | 204 | 57 | 38 | 28 | 128 | 455 |
| Only MCF-7 | 206 | 110 | 89 | 51 | 106 | 562 |
| Only MDA-MB-231 | 290 | 37 | 80 | 23 | 165 | 595 |
| MCF-10A and MCF-7 | 101 | 21 | 22 | 10 | 48 | 202 |
| MCF-10A and MDA-MB-231 | 129 | 11 | 27 | 13 | 93 | 273 |
| MCF-7 and MDA-MB-231 | 76 | 15 | 8 | 6 | 38 | 143 |
| All three | 231 | 65 | 35 | 27 | 144 | 502 |
| Total | 1237 | 316 | 299 | 158 | 722 | 2732 |

**Supplementary Table 3. Summary of structural variations detected in breast cell lines.** Structural variations types are deletions (DEL), translocations (TRA), duplications (DUP), inversions (INV), and insertions (INS), and are grouped by uniquely occurring (first three lines), commonly occurring in any combination of two cell lines (second three lines), and commonly occurring in all lines. SVs of < 1kb were filtered out.
